# Sex-specific role of epigenetic modification of a leptin upstream enhancer in the adipose tissue

**DOI:** 10.1101/2024.09.30.615808

**Authors:** Luise Müller, Rebecca Oelkrug, Jens Mittag, Anne Hoffmann, Adhideb Ghosh, Falko Noé, Christian Wolfrum, Esther Guiu Jurado, Nora Klöting, Arne Dietrich, Matthias Blüher, Peter Kovacs, Kerstin Krause, Maria Keller

## Abstract

**Objective:** Maternal hormonal status can have long-term effects on offspring metabolic health, likely regulated via epigenetic mechanisms. We elucidated the effect of maternal thyroid hormones on epigenetic regulation of *Leptin* (*Lep*) transcription in adipose tissue (AT) and investigated the role of DNA methylation at a *Lep* upstream enhancer (UE) in adipocyte biology.

**Methods:** Pregnant mice were treated with Triiodothyronine (T3). Offspring body composition and adipose tissue were analysed at 6 months (treatment: *N* = 8, control: *N* = 12). DNA methylation at the *Lep* UE and *Lep* mRNA levels in gonadal white adipose tissue (gWAT) and leptin serum levels were measured. The *Lep* UE was hypomethylated in murine preadipocytes using a dCas9-SunTag-TET1 system, followed by differentiation. In human visceral AT samples (*N* = 52) from the Leipzig Obesity BioBank (LOBB) *LEP* UE methylation and *LEP* mRNA levels were assessed alongside leptin serum levels and correlated with body fat percentage. Causal relationships were explored using mediation analysis.

**Results:** High maternal T3 levels reduced offspring body weight, total fat, and gWAT mass. Female offspring of T3-treated mothers showed lower *Lep* mRNA levels and higher *Lep* UE methylation compared to controls. CRISPR/dCas9 editing reduced *Lep* UE methylation by ∼20% in preadipocytes without altering *Lep* expression or lipid accumulation after differentiation. In humans, a mediation of sex-specific difference in body fat percentage via *Lep* UE methylation was found.

**Conclusion:** Maternal thyroid hormones might affect offspring gWAT *Lep* expression in a sex-specific manner, potentially driven by DNA methylation at the *Lep* UE, influencing body fat percentage.

## Introduction

Obesity is defined as an abnormal or excessive accumulation of body fat. In regard to preventing and treating obesity, it is important to understand the regulators of body weight and fat mass. One key master regulator of body weight is the hormone leptin, which is secreted primarily from white AT. This adipokine is crucial in energy homeostasis and metabolism playing a significant role in obesity [1,2]. Many studies of leptin function have focused on the hormone’s action in the brain via its receptors, while the regulatory elements that enable dynamic changes in *leptin* gene expression received less attention. *Leptin* transcription can be influenced by various factors such as insulin [3], hypoxia [4] and hormonal signals [5]. Moreover, a fat-specific long non-coding RNA (lncOb) was described, which is regulated alongside fat mass and influences *leptin* expression by interacting with redundant enhancers [6,7]. In mice, the absence of functional lncOb is linked to lower plasma leptin levels and increased adiposity, but these mice still respond to leptin treatment [6]. In humans, a single nucleotide polymorphism (rs10487505) in the corresponding region is associated with circulating leptin levels and obesity in a sex-specific manner [8], but not with *leptin* gene expression suggesting post-transcriptional mechanisms, e.g. by the regulation of long non-coding RNAs [9].

According to the developmental origins of health and disease (DOHaD) theory, the development of adipose tissue and the risk of developing obesity and metabolic diseases later in life are influenced by pre- and perinatal environmental exposures *in utero* [10–15]. Research has shown that maternal factors during pregnancy can significantly alter the hormonal environment of the developing foetus, leading to lasting effects on *Lep* regulation in offspring [13,16–20]. For instance, maternal high-fat diet exposure is linked to sustained increases in leptin levels and elevated blood pressure in offspring, mediated by epigenetic memory [20]. Maternal nutrition during pregnancy can modify the relationship between leptin levels, body fat, and caloric intake in offspring, leading to excess adiposity and metabolic issues in adulthood [17].

It is known that thyroid hormones are significant regulators of foetal tissue development and maturation [21]. Thyroid deficiency before birth alters adipose tissue development, leading to overgrowth of white adipocytes, disrupted thermogenesis, and changes in gene expression related to metabolism and insulin resistance, potentially increasing the risk of neonatal survival issues, obesity, and metabolic dysfunction later in life [22]. While the essential role of maternal thyroid hormones in foetal development is well established, the effects of maternal hyperthyroidism on offspring remain poorly understood. A recent study by Oelkrug et al. in mice demonstrated that maternal T3 treatment during gestation leads to improved glucose tolerance in adult male offspring and hyperactivity of brown adipose tissue thermogenesis in both sexes starting early after birth [23]. However, the precise mechanisms through which thyroid hormones influence foetal adipose tissue development remain unclear. Interestingly, elevated levels of choline, which is involved in the synthesis of S-adenosylmethionine, a major methyl donor required for DNA methylation, were found in the serum of the T3-treated dams [23]. This suggests that epigenetic mechanisms may play a role in mediating the effects of thyroid hormones on foetal adipose tissue development.

Epigenetic mechanisms, including DNA methylation, are believed to mediate the impact of prenatal stress and other factors on adiposity and obesity risk in children [11,24,25]. Pre- and perinatal environmental exposures, including diet and pollutants, can induce epigenetic changes that impact disease development in children [25]. Maternal intake of methyl-group donors can affect offspring health by altering DNA methylation patterns, linking early environmental exposure to disease risk in offspring [26]. Furthermore, Lecoutre et al. showed that maternal obesity might epigenetically program increased *Lep* gene expression via modulation of upstream enhancer DNA methylation in offspring, contributing to white adipose tissue accumulation [16]. This analysed enhancer is located around 36 kb upstream from the leptin gene and in close proximity to the long non-coding RNA lncOb.

Taken together, pre- and perinatal exposure to environmental factors can significantly influence the risk of developing obesity and metabolic diseases later in life. Among these factors, maternal thyroid hormone levels during pregnancy have emerged as a critical determinant of offspring metabolic health. Therefore, this study aims to elucidate the effects of maternal thyroid hormone levels on *Lep* expression in offspring, focusing on whether these effects are mediated through epigenetic regulation, specifically DNA methylation of the *leptin* upstream enhancer (*Lep* UE). Furthermore, this research explores whether epigenetic editing at this upstream enhancer can modulate *Lep* expression and adipogenesis *in vitro*. Finally, we examine the relationship between DNA methylation of the human *LEP* UE in OVAT, leptin expression, and body fat percentage.

## Material and Methods

### Animal model

This study analysed the effect of maternal hyperthyroidism on the offspring’s white adipose tissue function using a subset of samples from previously published mice [23]. Hyperthyroidism during pregnancy was induced in wild type female C57BL/6NCrl (Charles River, Germany) at the age of three to four month with 0.5□mg/L T3 (3,3’,5-Triiodo-L-thyronine, T6397, Sigma Aldrich, Germany) in drinking water with 0.01% BSA from day of positive plug check until gestational day 18 (= day before birth). Body composition of offspring mice was analysed using Minispec LF110 and Minispec Plus Software 6.0 (Bruker, USA) at the age of five to seven months. Serum was collected after two centrifugation steps at 4°C (1000 g) and stored at −20°C. Adipose tissue samples were collected at the day of sacrifice (male offspring: 5-6 months, female offspring: 6-7 months), weighted, snap frozen on dry ice, and stored at −80 °C until nucleic acid extraction. All animal experiments and procedures were approved by the Ministerium für Energiewende, Klimaschutz, Umwelt und Natur MEKUN Schleswig-Holstein, Germany.

### Human cohort

Human OVAT samples from the Leipzig Obesity BioBank (LOBB; https://www.helmholtz-munich.de/en/hi-mag/cohort/leipzig-obesity-bio-bank-lobb) were collected from 518 individuals with severe obesity (median BMI [IQR] = 48.1 [41.9-53.7] kg/m²) at the university hospital in Leipzig. For DNA methylation analysis of the *LEP* UE a subset of 52 individuals (female: *N* = 31, male: *N* = 21) with a median BMI [IQR] = 49.0 [41.5-52.6] kg/m² was used. OVAT samples were collected during elective laparoscopic abdominal surgery as previously described [27,28], immediately frozen, and stored at −80 °C. Exclusion criteria encompassed participants under 18 years of age, chronic substance, or alcohol misuse, smoking within the 12 months prior to surgery, acute inflammatory diseases, use of glitazones as concomitant medication, end-stage malignant diseases, weight loss exceeding 3% in the three months preceding surgery, uncontrolled thyroid disorder, and Cushing’s disease. All participants gave written informed consent before taking part in the study and have been informed of the purpose, risks and benefits of the biobank. The study was approved by the ethics committee of the University of Leipzig (#159-12-21052012).

Phenotyping of the LOBB includes assessment of age, sex (self-reported), BMI (kg/m²), body fat mass (%) detection using bioelectrical impedance analysis, and metabolic biochemical assessment. Leptin serum levels were determined by enzyme-linked immunosorbent assay (ELISA) (Leptin Human ELISA, Mediagnost, Germany) as previously described [8]. *Leptin* mRNA levels were extracted from RNA sequencing data, which is described in detail in the Suppl. Information. The phenotypic characterisation of the total cohort as well as for the subset for methylation analysis can be found in Table 1.

**Table 1.** Phenotypic characteristics of LOBB cohort.

|  | cohort | N | median [IQR] | female |  | male |  | female vs male |
| --- | --- | --- | --- | --- | --- | --- | --- | --- |
|  |  |  |  | N | median [IQR] | N | median [IQR] | P-value |
| BMI [kg/m<sup>2</sup>] | total | 518 | 48.1 [41.9-53.7] | 339 | 47.1 [41.8-52.7] | 179 | 50.0 [42.6-54.7] | 0.065 |
|  | methylation subset | 52 | 49.0 [41.5-52.6] | 31 | 46.5 [39.6-51.0] | 21 | 52.0 [47.7-57.6] | **0.031** |
| Age [y] | total | 518 | 50.4 [40.1-57.0] | 339 | 50.5 [40.8-57.0] | 179 | 50.1 [39.2-57.2] | 0.716 |
|  | methylation subset | 52 | 50.0 [39.3-56.8] | 31 | 54.0 [40.9-57.4] | 21 | 48.3 [35.2-53.5] | 0.376 |
| Body fat [%] | total | 511 | 48.8 [41.9-54.3] | 337 | 52.0 [47.0-57.1] | 174 | 41.8 [37.0-46.8] | **6.19E-33** |
|  | methylation subset | 52 | 47.3 [40.6-55.8] | 31 | 54.5 [42.0-58.7] | 21 | 44.8 [39.8-48.0] | **0.017** |
| TSH [mU/l] | total | 510 | 1.6 [1.0-2.2] | 335 | 1.5 [0.9-2.2] | 175 | 1.7 [1.1-2.2] | 0.072 |
|  | methylation subset | 52 | 1.9 [1.2-2.4] | 31 | 1.8 [1.2-2.4] | 21 | 1.9 [1.2-2.4] | 0.963 |
| fT3 [pmol/l] | total | 327 | 4.9 [4.3-5.4] | 212 | 4.8 [4.2-5.3] | 115 | 5.2 [4.5-5.6] | **0.002** |
|  | methylation subset | 46 | 5.0 [4.7-5.4] | 27 | 5.0 [4.6-5.3] | 19 | 5.2 [4.7-5.6] | 0.289 |
| fT4 [pmol/l] | total | 338 | 16.6 [14.6-18.9] | 222 | 17.0 [14.8-19.4] | 116 | 15.6 [14.5-18.1] | **0.011** |
|  | methylation subset | 48 | 17.0 [14.9-19.7] | 29 | 18.4 [15.8-21.6] | 19 | 15.8 [14.8-17.5] | 0.058 |
| LEP mRNA level [AU] | total | 518 | 12.2 [11.5-13.0] | 339 | 12.1 [11.4-12.9] | 179 | 12.5 [11.6-13.2] | **0.045** |
|  | methylation subset | 52 | 12.3 [11.4-13.0] | 31 | 12.3 [11.3-12.9] | 21 | 12.7 [11.5-13.9] | 0.274 |
| LEP serum level [ng/μl] | total | 485 | 40.2 [25.8-54.8] | 315 | 41.9 [30.4-57.8] | 170 | 31.2 [17.9-47.9] | **4.70E-07** |
|  | methylation subset | 50 | 41.6 [35.9-50.4] | 30 | 43.4 [36.4-51.3] | 20 | 41.1 [34.9-47.3] | 0.434 |
| LEP UE meth. [%]* | methylation subset | 52 | 63.5 [56.3-67.7] | 31 | 65.3 [60.8-68.5] | 21 | 59.1 [53.2-65.5] | **0.031** |

Data is shown as median with interquartile range [Q1-Q3]. Comparisons between female and male were calculated by unpaired Wilcoxon rank sum test. Significant P-values (< 0.05) were marked in bold.
AU - arbitrary units, BMI - Body Mass Index, TSH - thyroid-stimulating hormone, fT3 - free triiodothyronine, fT4 - free thyroxine, LEP - Leptin, UE - upstream enhancer
\* mean DNA methylation of 6 CpG sites

### Plasmid preparation for *in vitro* hypomethylation

The targeted *in vitro* hypomethylation was performed as described by Morita et al. using the pPlatTET-gRNA2 vector, a gift from Izuho Hatada (Addgene plasmid #82559; http://n2t.net/addgene:82559; RRID:Addgene_82559) [29]. This all-in-one plasmid contains the dCas9-SunTag and single-chain variable fragment (scFv)-TET1 catalytic domain (TET1CD) system, and a gRNA expression system (Figure 2A). Three different gRNAs targeting the murine *Lep* UE were designed using the CRISPR website from Dr. Feng Zhang’s laboratory (http://crispor.tefor.net) [30]. Cloning was performed by linearization of the pPlatTET-gRNA2 with Afl II (NEB, USA) and Gibson assembly-mediated incorporation of the gRNA insert fragment as described before [31]. The target sequences are listed in Suppl. Table 1.

### *In vitro Lep* UE hypomethylation

The workflow for *in vitro* hypomethylation procedure is illustrated in Figure 2B. We used SV40 T-antigen immortalised murine preadipocyte cells isolated from epididymal white adipose tissue of C57BL6 mice [32], which were grown at 37°C and 5% CO_2_ atmosphere in Dulbecco’s Modified Eagle’s Medium (DMEM)-high glucose (Gibco by Life Technologies, USA) supplemented with 10% Foetal Bovine Serum (FBS Superior S0615, Sigma Aldrich, Germany). Cells were transfected with in total 5µg of the all-in-one plasmids (1.67 µg for each gRNA) by electroporation using the Neon™ Transfection System (Invitrogen, Germany). For increased transfection efficiency of large-size CRISPR/Cas9 vectors [33], cells were co-transfected with the equimolar amount of an empty pGEM®-T vector (Promega, WI USA) with the size of around 3 kb. As negative control, cells were transfected with the pPlatTET-gRNA2 without incorporated gRNA (plasmid control). Cells were sorted 24 h post-transfection for GFP-positive cells using FACS Aria II (BD Biosciences) and the BD FACSDiva Software 9.0.1 (BD Biosciences) and seeded back to growth media supplemented with 1% Penicillin/Streptomycin to avoid potential contamination. The differentiation of cells is described in detail in Suppl. Information.

### Murine Leptin ELISA

Mouse leptin ELISA (Cat # 90030, CrystalChem, USA) was perfomed according to manufacture’s instruction. In brief, 5 µl of sample or standard were added per well and incubated overnight at 4°C. Optical density at 450/ 630 nm was measured using the BMG SPECTROstar Nano Microplate Reader.

### Bisulfite sequencing of human and murine *leptin* upstream enhancer

Genomic DNA extraction from murine gWAT, human OVAT, as well as from cultured adipocytes was performed as described in detail in Suppl. Information. DNA was bisulfite converted using the EpiTect Fast Bisulfite Conversion Kit (Qiagen, Germany) and further amplified using the EpiTect Whole Bisulfitome Kit (Qiagen, Germany). Target sequences were amplified using self-designed primers (Suppl. Table 1) and pyrosequencing was performed using the PyroMark Q24 technologies and corresponding Gold Kits (Qiagen, Germany). Non-template controls were included in all steps to rule out contaminations. All PCR products were quality controlled using agarose gel electrophoresis prior sequencing. Only methylation values of CpG positions with good quality (automatic blue calling) were considered for analysis. All experiments were performed in duplicates.

### Quantitative RT-PCR analysis

RNA extraction is described in detail in Suppl. Information. The cDNA synthesis from murine gWAT RNA was performed using QuantiTect Reverse Transcription Kit (Qiagen, Germany) with integrated removal of genomic DNA contamination and SuperScript III (Invitrogen) for RNA from cell culture samples. A non-enzyme control, as well as a non-target control without RNA was always carried along to control for genomic DNA and other contaminants. Quantitative real time PCR was performed using Power Up SYBR Green (Applied Biosystems) in the LightCycler 480 System (Roche). Self-designed primers can be found in Suppl.Table 1. Measurement of a standard curve with a serial dilution of pooled cDNA was included to determine efficiency of the PCR. Relative quantities of mRNA levels were determined using the efficiency based ΔCt method normalising to the mRNA level of *Rplp0* [34]. Measurements were performed in triplicates and averaged.

### Gene expression profiling of murine gWAT with Clariom S arrays

Gene expression profiling in the gWAT samples from the offspring of maternal T3 treatment experiment was performed by Core Facility for DNA technologies of Knut Krohn from the University of Leipzig using GeneChip Clariom S arrays (Affymetrix, Germany). The raw microarray data were preprocessed using the oligo R package (v1.50.0), which includes background correction and quantile normalisation through the Robust Multichip Average (RMA) algorithm [35,36]. Further quality control was carried out using both the Biobase (v2.46) and oligo R packages [37]. Genes with median transcript intensities below a threshold of 4 were filtered out from the normalised dataset. Differentially expressed genes (DEGs) were identified using the Linear Models for Microarray Data method in the R package limma (v3.42) [38], incorporating array weights to enhance the signal-to-noise ratio. The threshold for identification of DEGs was set as unadjusted *P* value < 0.01 and |log2 fold change (FC)| ≥ 0.5.

### Lipid staining and quantification using AdipoRed^TM^

Intracellular lipid accumulation was assessed by fluorescence spectroscopy after staining with AdipoRed™ reagent (Lonza, Switzerland). Successfully transfected cells were grown in 96-well plates and differentiated as described in Suppl. Information. At days 0, 4, and 8 of differentiation, intracellular lipids were stained with AdipoRed™ (1:40 in PBS, 10 min) and measured using the FLUOstar OPTIMA Microplate Reader (BMG Labtech GmbH, Germany; excitation 485 nm; emission 520 nm). DNA was then stained with Hoechst 33342 (Sigma Aldrich, Germany; 1:2000 in PBS, 15 min) and fluorescence intensity measured (excitation 355 nm; emission 460 nm) to estimate cell number. Relative lipid amount was calculated as the ratio of AdipoRed™ fluorescence intensity to Hoechst fluorescence intensity. With the same staining procedure microscopy of cells was performed using the Carl Zeiss Axio Observer Z1 (Carl Zeiss, Germany) with the filter set 38 HE green and ZEN software. Fluorescence images were captured using the FITC channel for lipids and DAPI channel for DNA.

### Statistics and bioinformatic analysis

Statistical analyses were performed using R v.4.4.1. Group comparisons of phenotypes (body weight, total fat mass, gWAT mass), leptin serum levels, *Lep* mRNA expression and DNA methylation of *Lep* UE between were performed using unpaired Wilcoxon rank sum test defining *P* < 0.05 as significant. Spearman’s rank correlation was performed to assess associations between continuous variables using *ggcormat* from the R package *ggstatsplot* v.0.12.4. Effects of *in vitro* hypomethylation treatment on *Lep* UE DNA methylation, *Lep* mRNA level and lipid accumulation during course of adipocyte differentiation was analysed by two-way mixed ANOVA, using time as within factor and treatment as between factor. Simple main effects were further analysed by pairwise paired Student’s t test. Overrepresentation analysis for Wikipathways with the gene lists, which were found to be differentially expressed between offspring gWAT of T3 treated vs control dams, was performed using clusterProfiler v.4.12.2 [39]. The analysis was done separately for upregulated (log2 FC > 0, *P* < 0.05) and downregulated genes (log2 FC < 0, *P* < 0.05). Causal mediation analysis was conducted to address two assumptions: i) if the observed sex-specific differences in body fat percentage are mediated via LEP UE DNA methylation in human OVAT and ii) if the relationship between the female specific LEP UE DNA methylation and body fat percentage is mediated by leptin serum levels. The mediation analyses were performed using *mediation* R package v.4.5.0. We employed bootstrapping procedures to test the significance of the indirect effects, i.e. we generated 1,000 bootstrapped samples to estimate the unstandardized indirect effects. The 95% confidence intervals for the indirect effects were computed by determining the 2.5th and 97.5th percentiles of the bootstrapped indirect effects.

## Results

### Offspring’s body composition after maternal T3 treatment

High maternal T3 levels during pregnancy led to a significantly reduced body weight [g], total fat mass and gWAT mass relative to body weight [%] in the adult offspring (Figure 1B-D, *P* < 0.02). When grouped by sex, only body weight was significantly reduced in female offspring (*P* = 0.007, Suppl. Figure 1A-C). Apart from this, serum levels of thyroid hormones (T3, T4, and TSH) were not altered in offspring of T3 treated dams compared to control offspring [23] in agreement with similar previous studies [40,41].

**Figure 1.**
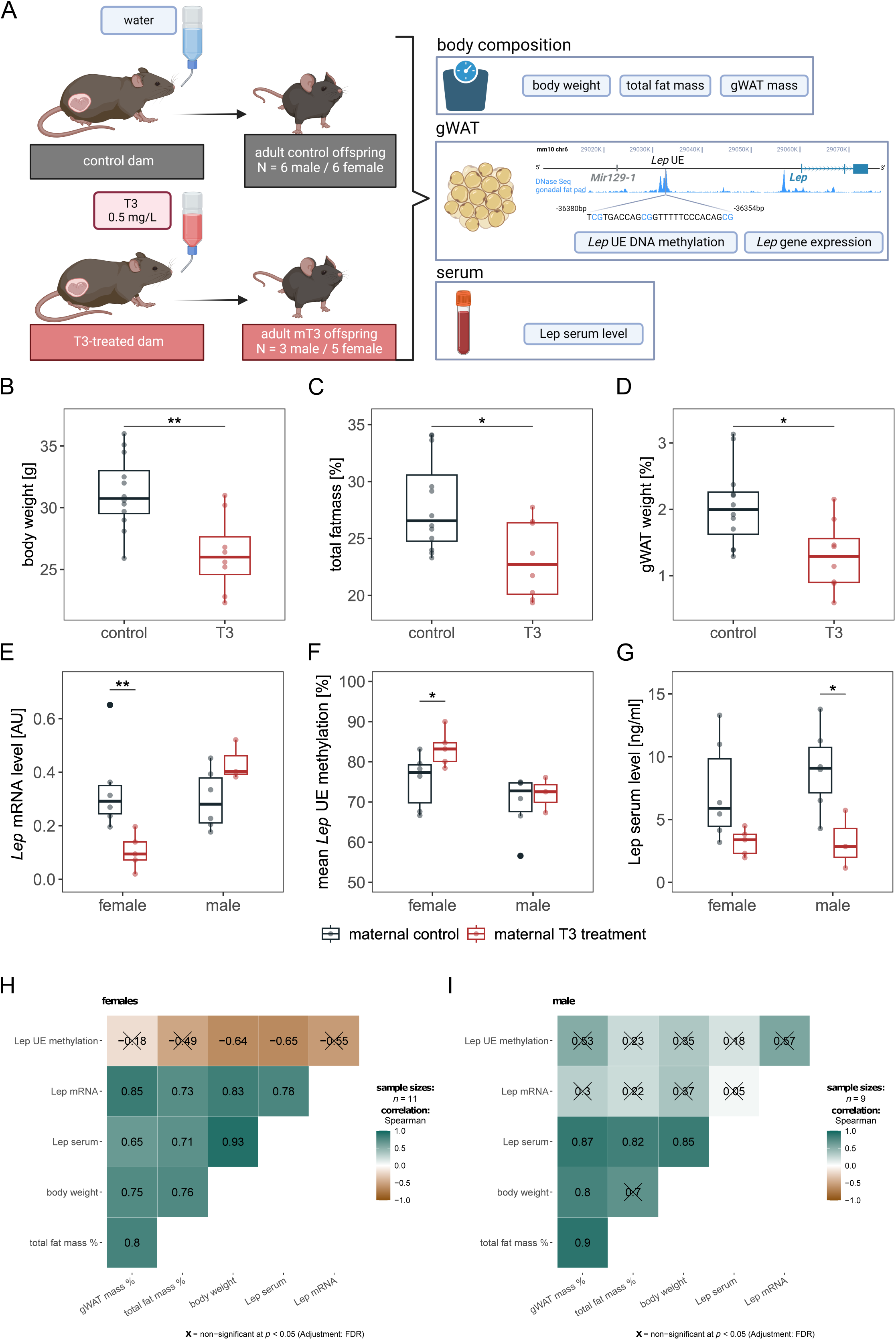
Effects of high maternal thyroid hormone during gestation on murine adipose tissue biology in offspring and *Leptin* mRNA expression. **(A)** Scheme of experimental design. Genomic region plot was adapted from epigenome browser (http://epigenomegateway.wustl.edu/browser/) including information about DNase sequencing data of gonadal white adipose tissue (gWAT). Location of the analysed target sequence including three CpG sites in the analysed *Lep* enhancer (∼ 36 kb upstream of the transcription start site of *Lep)* is shown. Scheme was generated using biorender.com. **B-G)** Boxplots show alterations in **B)** body weight [g], **C)** total fat mass [% body weight], **D)** gonadal fat mass [% body weight] and **E)** *Lep* mRNA level in gWAT analysed by quantitative real time PCR and normalised to the level of *Rplp0*, **F)** average DNA methylation level [%] of *Lep* upstream enhancer across all three analysed CpG sites by bisulfite pyrosequencing in gWAT from offspring, and **G)** Leptin serum level (ng/ml) determined by ELISA for offspring of T3 treated dams (in red, *N* = 5 female, *N* = 3 male) compared to offspring from control dams (in gray, *N* = 6 female, *N* = 6 male). Significance levels of differences calculated by unpaired Wilcoxon rank sum test are shown for *** *P* < 0.001, ** *P* < 0.01, * *P* < 0.05. **H/I)** Heatmaps show spearman correlation coefficients of enhancer DNA methylation, mRNA level and serum levels of leptin with anthropometric phenotypes for **H)** female offspring (mT3: *N* = 5, control: *N* = 6) and **I)** male offspring (mT3: *N* = 3, control: *N* = 6). Positive correlations are shown in petrol and negative correlations in brown. Non-significant correlations (FDR < 0.05) are crossed out.

### Leptin expression and methylation levels in offspring after maternal T3 treatment

We observed a significant interaction between sex and the maternal T3 treatment on *Lep* mRNA level in offspring gWAT (F(1,16) = 11.179, *P* = 0.004, eta2[g] = 0.411) suggesting a sex-specific effect of maternal T3 treatment on *Lep* mRNA level in the offspring. While significantly lower *Lep* mRNA level in female offspring of T3-treated mothers compared to control offspring could be observed (*P* = 0.009, Figure 1E), *Lep* mRNA levels in gWAT of male offspring born to T3 treated dams were, albeit higher, not significantly different (*P* = 0.167, Figure 1E) from that of control dams.

The gWAT DNA methylation at the previously reported enhancer region ∼ 36 kb upstream of *Lep* revealed no significant interaction effect of sex and T3 treatment. However, the DNA methylation was significantly higher in female offspring of T3 treated mothers (Figure 1F, median methylation difference = 5.9 %, *P* = 0.03) compared to control animals. No differential methylation was observed between male offspring born to T3 treated- and control dams (Figure 1F, median methylation difference = 0.3 %, *P* = 0.714). On leptin serum levels in offspring no significant interaction effect of sex and T3 treatment was found, but an effect of T3 treatment was evident (F(1,16) = 11.274, *P* = 0.004, eta2[g] = 0.413). In total, maternal T3 treatment was related to a significantly lower leptin serum level in offspring (*P* = 0.002). After grouping by sex the difference was only significant in male offspring (*P* = 0.034, Figure 1G), but in female offspring of T3 treated mothers still a trend of lower leptin serum levels could be observed (*P* = 0.058, Figure 1G) when compared to controls.

Irrespective of maternal treatment, we observed in female offspring a strong positive correlation of *Lep* mRNA expression in gWAT as well as leptin serum levels with gWAT mass (% of body weight), total fat mass (% of body weight) and body weight (Figure 1H, FDR < 0.05, Spearman rho > 0.6). In male offspring we observed this positive correlation only for leptin serum levels with gWAT mass (% of body weight), total fat mass (% of body weight) and body weight (Figure 1I, FDR < 0.05, Spearman rho > 0.8), while *Lep* mRNA expression in gWAT does not correlate with any trait. In female offspring the *Lep* UE methylation correlates negatively with body weight and leptin serum level (Figure 1H, FDR < 0.05, Spearman rho < −0.6), but not significantly with *Lep* mRNA expression in female offspring. In male offspring the *Lep* UE DNA methylation in gWAT does not correlate with any trait at all (Figure 1I, FDR > 0.05).

### Genome-wide expression changes in offspring’s gWAT after maternal T3 treatment

To further explore molecular determinants of altered body composition in offspring of T3 treated vs control dams we performed microarray gene expression analysis in gWAT of adult offspring. While the effect sizes of maternal T3 treatment on the adult offspring might not be sufficient to detect genome-wide significant differences in gene expression after correction for multiple testing, we only identified several DEGs using relaxed cut offs with |log2 FC| > 0.5 and an unadjusted *P* < 0.01 (females: 291 DEGs, Suppl. Table 2; males: 862 DEGs, Suppl. Table 3). A summary of the most significant DEGs can be found in Volcano plots (Suppl. Figure 2). Notably, *Lep* was among the most significantly altered genes in female offspring (log2 FC = −0.84, *P* = 0.0026), consistent with our real-time PCR results. To gain further insights, we performed a Wikipathway overrepresentation analysis on upregulated and downregulated genes in adult offspring following maternal T3 treatment. The analysis in the female offspring revealed that upregulated genes (*Creb1*, *Foxo1*, *Gata2*, *Nr2f2*, *Rora*, *Klf5)* are involved in the ‘white fat cell differentiation’ (*WP2872*, FDR = 0.048; Suppl. Figure 3A, Suppl. Table 4). Additionally, six (*Mvk*, *Dhcr7*, *Lss*, *Idi1*, *Hmgcr*, *Sqle*) from the fifteen genes of the ‘cholesterol biosynthesis’ pathway (*WP103*, FDR = 0.001) were among the downregulated genes in female offspring of T3 treated dams compared to control animals (Suppl. Figure 3A, Suppl. Table 4). In gWAT of male offspring this analysis revealed that gene expression of 20 out of 268 ‘non-odorant G-Protein coupled receptors‘(GPCRs) is upregulated (*WP1396*, FDR = 0.035), while 16 genes from 81 ‘cytoplasmic ribosomal proteins’ (*WP 163*, FDR = 2 × 10^-6^) were downregulated in offspring of T3 treated compared to control dams (Suppl. Figure 3B, Suppl. Table 5).

### *In vitro* hypomethylation of *Lep* upstream enhancer using a dCas9-Suntag-TET1 system

With the aim to investigate the functional role of DNA methylation at the *Leptin* upstream enhancer *in vitro,* the target region was hypomethylated in immortalised murine epididymal preadipocytes using a system consisting of a dCas9–SunTag and scFv–TET1 catalytic domain (Figure 2A). The workflow of the experiment is illustrated in Figure 2B. After transfection we achieved an approximately 20 % reduction of *Lep* UE DNA methylation in successfully transfected cells (mean DNA methylation ± SD = 71.0 ± 4.6 %) compared to untreated cells (mean DNA methylation ± SD = 90.4 ± 6.4 %) at day of induction (= day 0). We further observed no significant increase in DNA methylation during adipocyte differentiation course in the transfected cells (Suppl. Figure 4) indicating a stable reduction of DNA methylation during differentiation. After combining DNA methylation of all time points we observed a significantly different DNA methylation comparing the treatments (Figure 2C, *P* = 5.1 × 10^-4^) and pairwise comparisons confirm a significantly lower DNA methylation in the cells transfected with the vector targeting the *Lep* UE (median DNA methylation [IQR] = 73.6 [71.4 - 74.3] %) compared to untreated cells (median DNA methylation [IQR] = 91.4 [83.8 - 96.2] %, FDR < 0.001) and cells transfected with the plasmid control without gRNA (median DNA methylation [IQR] = 88.5 [75.9 - 94.1] %, FDR < 0.001). We observed no significantly different DNA methylation in the cells transfected with the plasmid control against the untreated cells (FDR = 0.87) indicating no significant global effect due to increased amounts of TET1 catalytic domain per se.

**Figure 2.**
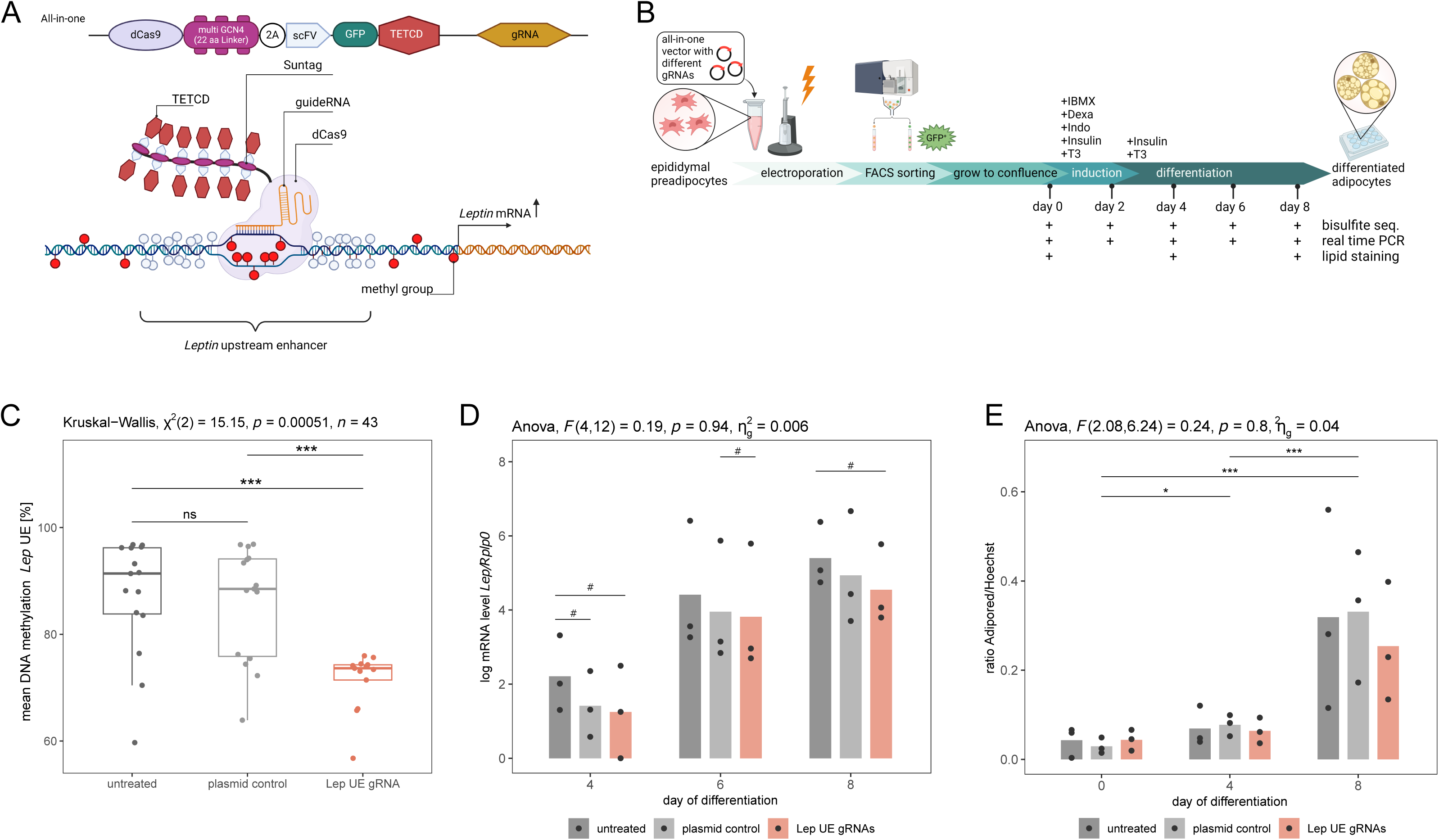
*In vitro Lep* UE hypomethylation and effect on adipocyte differentiation. **A)** Principal scheme of targeted *in vitro* hypomethylation using a dCas9-Suntag-TET1 system showing in the top the “all in one” vector which contains dCas9 peptide array (linker length: 22aa), antibody-sfGFP-TET1CD, and gRNA expression system. Scheme was generated using biorender.com **B)** Workflow of the experiment, which is described in detail in the methods section. For the hypomethylation epididymal preadipocytes were transfected by electroporation with all-in-one vectors incl. three different gRNAs targeting the *Lep* UE (= Lep UE gRNA, red). For control we used untreated cells (= UT, dark grey) and cells transfected with the all-in-one vector without gRNA (= plasmid control, light grey). Workflow was generated using biorender.com **C)** Boxplot shows DNA methylation level [%] of *Lep* UE across all three analysed CpG sites by bisulfite pyrosequencing. Significance of differences between treatments were calculated using Kruskal Wallis test, followed by pairwise comparisons by Wilcoxon rank sum test corrected for multiple testing by FDR (*** FDR > 0.001, ns FDR > 0.05). Dots represent mean methylation levels of day 0, 2, 4, 6 and 8 of differentiation for each experiment (n = 3 experiments x 5 time points). **D)** Barplot shows the mean of log transformed *Lep* mRNA levels normalised to *Rplp0* mRNA levels for days 4, 6 and 8 of adipocyte differentiation from n = 3 experiments. Results of mixed two-way ANOVA used to assess the effect of treatment and time on *Lep* expression are shown on top of the graph. Pairwise comparisons of treatment effects using paired Student’s t-test (paired by experiment number) are depicted in the graph. No significant differences were found after correction for multiple testing by FDR. Shown significance level are uncorrected for multiple testing: # *P* < 0.05. **E)** Barplot shows the lipid amounts in cells during differentiation on day 0, 4 and 8 between treatments. Shown is the average ratio of Adipored and Hoechst fluorescence intensity from n = 12 wells in n = 3 experiments. Results of mixed two-way ANOVA used to assess the effect of treatment and time on lipid accumulation are shown on top of the graph. Significance levels from pairwise comparison of time points using paired Student’s t-test (paired by experiment number) and corrected for multiple testing by FDR are depicted in the graph (*** FDR < 0.001, ** FDR < 0.01, * FDR < 0.05). No significant differences are found when comparing treatments.

### Effect of upstream enhancer hypomethylation on *Lep* mRNA level during adipocyte differentiation in epididymal preadipocytes

Effects of UE hypomethylation on the *Lep* mRNA level and lipid accumulation during differentiation were evaluated. Since *Lep* mRNA levels were not detectable at the day of induction and two days later, only mRNA levels of day 4 and later were considered. No significant interaction of treatment and time on the *Lep* mRNA level was found (Anova, F(4,12) = 0.19, *P* = 0.94, Figure 2D). A simple main effect of time on the *Lep* mRNA level was found proving that *Lep* mRNA level was rising in the course of adipocyte differentiation (Anova, F(2,12) = 110.119, *P* = 1.9 × 10^-8^). We did not observe a significant effect of treatment on *Lep* mRNA level (Anova, F(2,12) = 0.309, *P* = 0.745). After pairwise comparisons we observed nominal lower *Lep* mRNA levels in transfected cells compared to untreated cells at day 4 and day 8 (*P* < 0.05, Figure 2D). However, this trend of lower *Lep* expression was also observed in the cells transfected with the plasmid control compared to untreated cells (*P* < 0.05). At day 6 we observed a slightly lower *Lep* expression in treated cells (Lep UE gRNAs) compared to cells transfected with the plasmid control (*P* < 0.05). The observed differences were moderate and none of these comparisons was statistically significantly different after correction for multiple testing.

### Effect of upstream enhancer hypomethylation on lipid accumulation during adipocyte differentiation in epididymal preadipocytes

The effect of *Lep* UE hypomethylation on the lipid accumulation during adipocyte differentiation was assessed by AdipoRed staining of lipids. No significant interaction was observed between hypomethylation effect and time on the lipid accumulation (Anova, F(2.08, 6.24) = 0.24, *P* = 0.8, Figure 2E). A significant increase of lipids during the course of adipocyte differentiation (Anova, F(1.04,6.24) = 23.998, *P* = 0.002) was confirmed by pairwise comparisons (FDR < 0.05, Figure 2E) and supported the accumulation of lipids during the differentiation. However, no significant difference was found in cells after hypomethylation of the *Lep* UE compared to the control cells (Figure 2E).

### Effect of *in vitro* hypomethylation of *Lep* UE in other adipocyte cell lines

Additionally, we performed the experiment in 3T3L1 cells as well as in immortalised female cells from the inguinal (subcutaneous) fat depot. In 3T3L1 cells we achieved on average a 35% hypomethylation of the *Lep* UE in transfected vs. untreated cells (Suppl. Figure 5A). In these cells we observed at day 4 a significant difference of *Lep* mRNA level between hypomethylated and untreated cells (FDR < 0.01, Suppl. Figure 5B), but not when comparing with the plasmid control cells and this difference was not seen in the following days of differentiation. Also, in 3T3L1 cells we observed no effect of *Lep* UE hypomethylation on lipid accumulation (Suppl. Figure 5C). In the inguinal cells from female mice, we achieved on average 21 % hypomethylation of *Lep* UE (transfected vs. untreated inguinal cells, Suppl. Figure 6A) taking all time points together. Although we were not able to detect *Lep* mRNA expression despite successful adipocyte differentiation, we observed a trend of reduced lipid accumulation after hypomethylation of *Lep* UE in these cells (Suppl. Figure 6B/C).

### Sex-specific differences in leptin expression and methylation levels in human visceral adipose tissue

Bisulfite sequencing of a corresponding *LEP* UE region in human OVAT (*N* = 52) revealed significant higher methylation levels in female (*N* = 31, median methylation [IQR] = 65.3 [60.8 - 68.5] %) compared to male individuals (*N* = 21, median methylation [IQR] = 59.1 [53.2 - 65.5] %, *P* = 0.031, Table 1). In the entire cohort, in female subjects we observed a significantly lower OVAT *LEP* mRNA expression (*N* = 518, *P* = 0.045, Table 1), while Leptin serum levels were significantly higher (*N* = 485, *P* < 0.0001, Table 1) than in males.

### Sex-specific causal relationship of human *LEP* upstream enhancer methylation with body fat percentage

Correlation analysis with DNA methylation at the *LEP* upstream enhancer revealed a significant positive correlation with body fat (% of body weight) in female individuals with obesity (Figure 3A, Spearman rho = 0.61, FDR < 0.05). We also observed a positive correlation of *LEP* methylation with LEP serum levels in female individuals (Figure 3A, Spearman rho = 0.38, FDR < 0.1). We found no significant association of *LEP* UE methylation with *LEP* mRNA level (FDR > 0.1). In male individuals with obesity the *LEP* UE methylation no correlation with any trait was identified (Figure 3B, FDR > 0.1). Also, *LEP* mRNA level in OVAT and LEP serum levels correlated with no other trait in males. Since *LEP* UE methylation did not correlate with *LEP* mRNA expression in OVAT, we addressed another causal relationship of DNA methylation with body fat percentage. First, we tested in a mediation analysis if *LEP* UE DNA methylation functions as a mediator for the sex-specific difference in body fat percentage (Figure 3C, Table 2). In our model, the sex-specific effect on body fat percentage was fully mediated via the DNA methylation in three out of six analysed CpG sites (Table 2, CpG 4-6, ADE *P* > 0.1, ACME *P* < 0.05) and partially mediated for CpG site 1 (ADE *P* < 0.1, ACME *P* < 0.05). The proportion of mediation ranged from 30.8% (CpG site 1) to 48.1 % (CpG site 4). Since the *LEP* UE methylation correlated only in females with the body fat percentage, we further used a second mediation analysis to test if this association is mediated via the Leptin serum level in females (Figure 3D, Table 3). The second mediation analysis revealed for the DNA methylation at CpG site 1, 4 and 6 as well as for the mean DNA methylation over all CpG sites that the effect on the body fat percentage was partially mediated via the Leptin serum level (Table 3, ADE *P* < 0.01, ACME *P* < 0.05). The proportion of the mediation effect ranged from 27.6 % (CpG site 6) to 32.5 % (CpG site 4).

**Figure 3.**
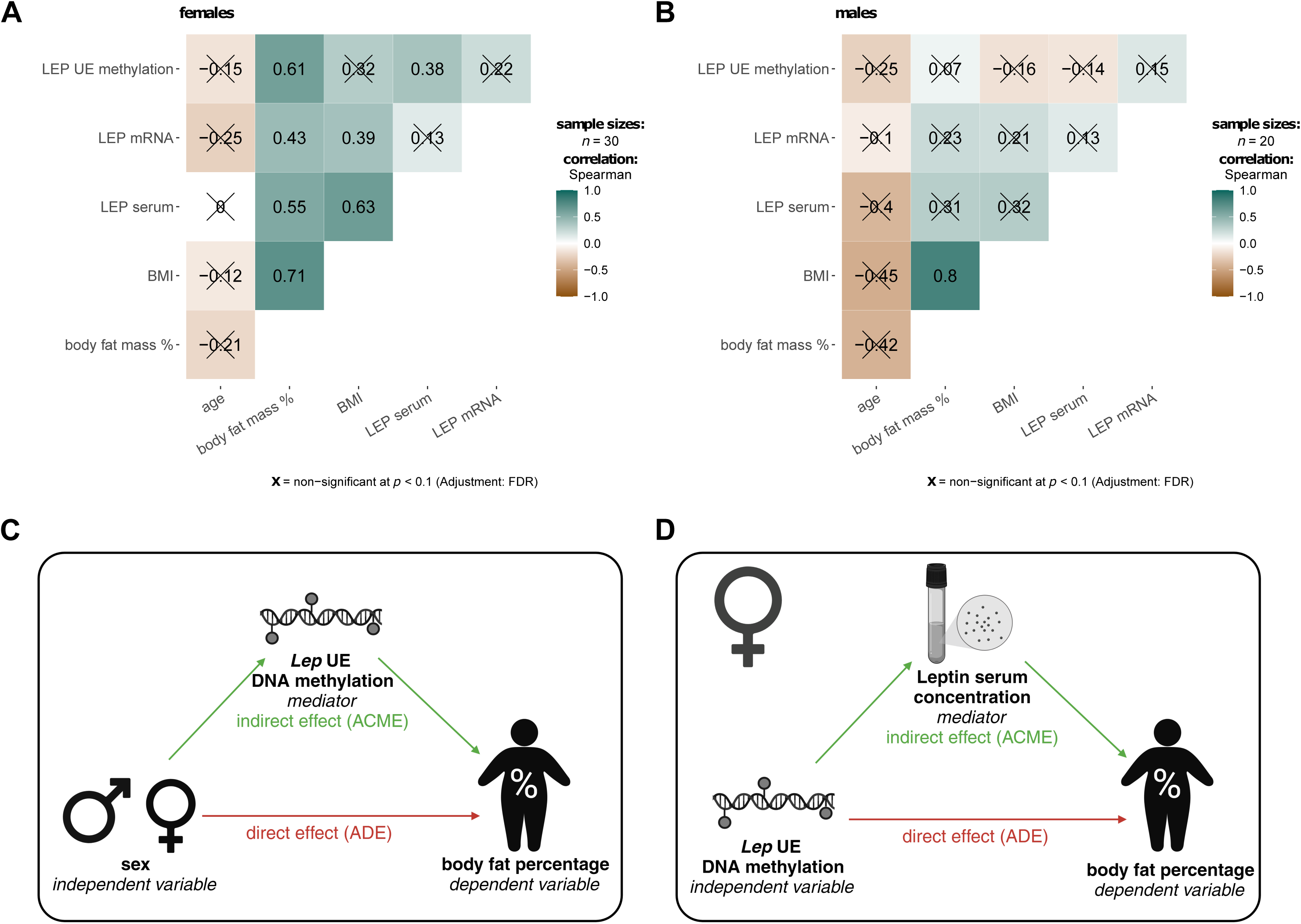
Relationship between sex-specific differences in leptin expression and enhancer DNA methylation levels in human visceral adipose tissue with body fat percentage. Heatmaps show spearman correlation coefficients of enhancer DNA methylation, mRNA level and serum levels of leptin with anthropometric phenotypes for female (**A**) and male (**B**) individuals with obesity. Positive correlations are shown in petrol and negative correlations in brown. Non-significant correlations (FDR < 0.1) are crossed out. C) Scheme for causal mediation analysis exploring the mediation of sex-specific differences in body fat percentage via DNA methylation at the *LEP* upstream enhancer. Results of the mediation analysis are shown in Table 2. D) Scheme for causal mediation analysis in female individuals exploring the mediation of the effect of DNA methylation at the *LEP* upstream enhancer on body fat percentage via leptin serum level. Results of the analysis are shown in Table 3.

**Table 2.**
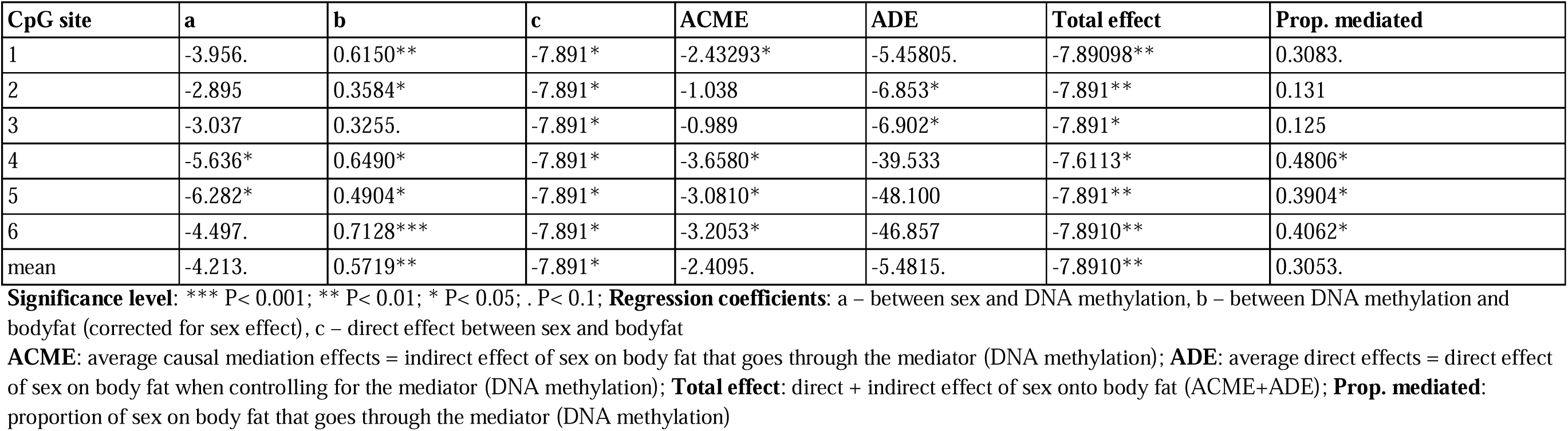
Causal mediation analysis of sex > LEP UE methylation > body fat percentage.

**Table 3.**
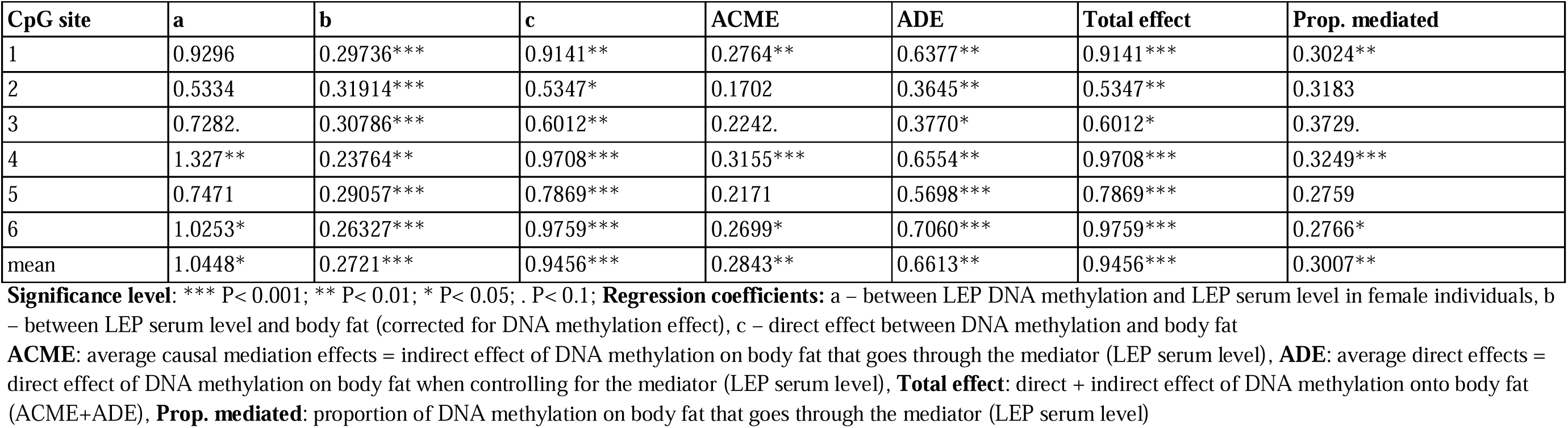
Causal mediation analysis of LEP UE methylation > LEP serum level > body fat percentage (in female individuals)

| CpG site | a | b | c | ACME | ADE | Total effect | Prop. mediated |
| --- | --- | --- | --- | --- | --- | --- | --- |
| 1 | 0.9296 | 0.29736*** | 0.9141** | 0.2764** | 0.6377** | 0.9141*** | 0.3024** |
| 2 | 0.5334 | 0.31914*** | 0.5347* | 0.1702 | 0.3645** | 0.5347** | 0.3183 |
| 3 | 0.7282. | 0.30786*** | 0.6012** | 0.2242. | 0.3770* | 0.6012* | 0.3729. |
| 4 | 1.327** | 0.23764** | 0.9708*** | 0.3155*** | 0.6554** | 0.9708*** | 0.3249*** |
| 5 | 0.7471 | 0.29057*** | 0.7869*** | 0.2171 | 0.5698*** | 0.7869*** | 0.2759 |
| 6 | 1.0253* | 0.26327*** | 0.9759*** | 0.2699* | 0.7060*** | 0.9759*** | 0.2766* |
| mean | 1.0448* | 0.2721*** | 0.9456*** | 0.2843** | 0.6613** | 0.9456*** | 0.3007** |

**Significance level:** \*\*\* P< 0.001; \*\* P< 0.01; \* P< 0.05; . P< 0.1; **Regression coefficients:** a – between LEP DNA methylation and LEP serum level in female individuals, b – between LEP serum level and body fat (corrected for DNA methylation effect), c – direct effect between DNA methylation and body fat
**ACME:** average causal mediation effects = indirect effect of DNA methylation on body fat that goes through the mediator (LEP serum level), **ADE:** average direct effects = direct effect of DNA methylation on body fat when controlling for the mediator (LEP serum level), **Total effect:** direct + indirect effect of DNA methylation onto body fat (ACME+ADE), **Prop. mediated:** proportion of DNA methylation on body fat that goes through the mediator (LEP serum level)

## Discussion

This study provides novel insights into the possible role of maternal thyroid hormones in shaping offspring’s metabolic outcome by analysing DNA methylation and gene expression in gWAT of adult offspring after maternal T3 treatment during pregnancy. Our findings underscore the significance of maternal thyroid hormones during pregnancy in influencing adult offspring’s body composition and gives insights into the sex-specific influence of DNA methylation at the *Lep* upstream enhancer in leptin expression and body fat percentage.

In the present study, maternal T3 treatment resulted in reduced body weight, total fat mass, and gWAT mass in adult offspring of both sexes. The increased DNA methylation at the *Lep* UE and the corresponding decrease in *Lep* mRNA expression in female offspring indicated that high T3 exposure *in utero* might epigenetically program *Lep* expression, contributing to altered adipose tissue function and reduced adiposity. In contrast, male offspring did not exhibit significant changes in *Lep* UE methylation but did show altered leptin serum levels, suggesting that the mechanisms by which maternal T3 influences metabolic outcomes may differ between sexes. One possible mechanism involves the differential expression and activity of thyroid hormone receptors and enzymes in the placenta and foetal tissues. The placenta mediates the metabolism of thyroid hormones, converting T4 to T3, and this process can vary between male and female foetuses. Studies indicate that the placental deiodinase, which regulates T3 availability, is differentially expressed based on foetal sex, potentially leading to variations in metabolic programming [42].

The genome-wide expression analysis further revealed several differentially expressed genes between offspring gWAT of T3 treated and control dams, with notable involvement in pathways related to adipogenesis and lipid metabolism. In females, genes associated with white fat cell differentiation were upregulated, while genes involved in cholesterol biosynthesis were downregulated in maternal T3 offspring. Since four (*Foxo1, Gata2, Nr2f2, Rora*) out of 6 upregulated genes related to white fat cell differentiation are actually inhibiting differentiation, one may assume that adipogenesis and lipogenesis might be inhibited in gWAT of female offspring of T3 treated dams. In males, the upregulation of non-odorant GPCRs might indicate alterations in signalling pathways that could influence energy homeostasis. GPCRs are implicated in the regulation of energy expenditure through mechanisms such as thermogenesis. Recent studies have explored the potential of pharmacological agents that target GPCRs to induce browning of white adipose tissue and enhance energy expenditure, which could be beneficial in combating obesity [43]. The different pathways targeted by these genes between female and male offspring after maternal T3 treatment further suggest sex-specific diversity in hormonal influences *in utero*.

The *in vitro* hypomethylation of the *Lep* UE in epididymal murine preadipocytes did not result in significant changes in *Lep* gene expression, which could be attributed to several factors. The reduction in methylation in this region alone might not be sufficient to regulate gene expression. Research has shown that even when methylation is reduced, other regulatory mechanisms, such as chromatin state or transcription factor availability, can maintain gene expression unaffected [44]. It is also possible that DNA methylation in this region affects transcription of other genes, such as the long noncoding RNA (lncOb) described by Dallner et al., which might regulate post-transcriptional events rather than directly influencing *Lep* gene expression [6]. Moreover, Lo et al. showed, that lncOb knock-down led to impaired adipogenesis, whereas overexpression of lncOb did not significantly alter leptin levels or other adipocyte markers suggesting that lncOb appears necessary for proper leptin regulation and adipogenesis, but alone is not sufficient to drive these processes [7].

Sex-specific regulations might also be the reason for absence of a detectable effect of the DNA hypomethylation at this region, since we used male murine cells. However, the relationship of DNA methylation at this region with leptin expression or body fat regulation was mainly observed in females. The experiment in female inguinal cells suggested that hypomethylation of the upstream enhancer region affected lipid accumulation, indicating a possible female-specific regulation. Sex-specific differences of *in vitro* leptin production and secretion in response to oestrogen were already shown for adipocytes from female vs. male sources [45]. However, since *Lep* expression was not detectable in the female inguinal cells, the mechanistic explanation for the role of DNA methylation in lipid accumulation remains unclear.

Furthermore, it is possible that the *in vitro* cell culture conditions were not conducive to cultivating leptin expressing adipocytes. Evidence suggests the existence of different types of adipocytes, each characterised by distinct gene expression patterns, with only certain adipocytes expressing leptin [46]. Moreover, adipocytes are influenced *in vivo* by various cell types such as macrophages, endothelial cells, fibroblasts, and neurons. These cells release cytokines, growth factors, and hormones that regulate adipocyte function, including leptin expression. *In vitro*, this network of cellular crosstalk is missing or incomplete. Therefore, *in vitro* models can mimic some aspects of adipocyte biology, but they often fall short in fully replicating the conditions needed for proper leptin expression. To further explore the role of this enhancer region in the regulation of leptin expression it might be necessary to use a cell culture system providing interaction of different cell types, such as spheroids, or to analyse the epigenetic modification using *in vivo* models.

Our data demonstrates a significant sex specific difference in *LEP* UE DNA methylation in OVAT from individuals with severe obesity with females exhibiting higher DNA methylation levels compared to males. Moreover, female subjects exhibit lower mRNA expression levels, but higher leptin serum levels than males. The higher serum levels of leptin in women are well-documented and are attributed to greater proportion of adipose tissue, but also to a higher production rate of leptin per unit mass of adipose tissue [47–49]. Furthermore, the influence of sex hormones, particularly oestrogens, has been suggested to modulate leptin expression, although the exact mechanisms remain complex and not fully understood [50,51]. In terms of fat depot specificity, leptin expression varies significantly across different adipose tissue depots. Studies have shown that *LEP* mRNA levels are generally higher in subcutaneous fat compared to visceral fat depots, such as omental fat [52–54]. The higher mRNA expression of *LEP* in OVAT of men despite lower serum levels might be due to lower AT mass or a higher contribution of the subcutaneous AT to leptin serum levels.

Despite missing correlations between *LEP* UE DNA methylation and *LEP* mRNA expression in human OVAT, we identified correlations with body fat percentage and leptin serum levels, exclusively in females. Using mediation analysis, we could show a significant mediation of the sex-specific variation in body fat percentage via the *LEP* UE DNA methylation. Additionally, a second analysis in females could show that the correlation between *LEP* UE DNA methylation and body fat percentage may be partially mediated by leptin serum levels. The mechanism by which DNA methylation affects body fat percentage and leptin serum levels, despite the absence of a significant effect on gene expression, remains unclear. However, post-transcriptional regulation of leptin expression via long non-coding RNAs could explain this observation. The reported leptin-lowering effect of the lncOb rs10487505 polymorphism, without any change in *LEP* mRNA expression [8], supports this hypothesis.

Despite the novel molecular insights into the role of maternal thyroid hormones in shaping the metabolic outcomes of offspring, several limitations of the present study must be acknowledged. The small sample sizes limit the generalizability of our findings. Larger cohorts would be necessary to confirm the observed sex-specific differences in DNA methylation and gene expression related to LEP and further elucidate the relationship between maternal T3 exposure and offspring metabolic outcomes. Furthermore, the lack of leptin-expressing adipocytes in *in vitro* experiments could have influenced our ability to detect significant changes in *Lep* gene expression following DNA methylation manipulation. Given that *Lep* gene expression is restricted to certain adipocyte subtypes, it is possible that the culture conditions were not conducive to maintaining or differentiating leptin-expressing cells, which might have contributed to the observed lack of response. Moreover, the human samples used in the study were derived from individuals with obesity, a condition often associated with leptin resistance. This resistance might have obscured potential correlations between *LEP* mRNA expression and leptin serum levels. Moreover, the metabolic profile of individuals with obesity may not accurately reflect broader population trends, limiting the applicability of these findings to non-obese individuals. Lastly, while the animal model allowed for a controlled investigation of maternal T3 effects, significant differences exist between murine and human physiology. For instance, species-specific variations in adipose tissue function, hormone regulation, and fat depot distribution may have impacted the comparability of the results. These interspecies differences could limit the direct translation of findings from the animal model to human populations.

## Conclusion

In summary, our data indicate the influence of maternal thyroid hormones on offspring’s gWAT *Lep* transcription in a sex-specific manner, potentially related to DNA methylation changes in the *Lep* upstream enhancer. Our data points towards a causal relationship of a sex-specific DNA methylation effect at the human *LEP* upstream enhancer on the body fat percentage, mediated by leptin serum levels in females. The success of targeted epigenetic editing might promote the use of similar approaches for novel strategies in treatment of obesity and related metabolic disorders.

## Author contributions

Conceptualization: JM, KK, MK; Methodology: LM, RO, AH; Formal analysis: LM, AH; Investigation: LM, RO; Resources: RO, JM, NK, AD, MB; Data curation: AH, AG, FN, CW; Writing original draft: LM, MK; Writing – Review and Editing: LM, RO, AH, EG, PK, KK; Visualisation: LM; Supervision: PK, KK, MK; Project coordination: KK, MK; Funding acquisition: JM, MB, PK, KK, MK

## Acknowledgement

Immortalised epididymal preadipocytes were kindly provided by Prof. Klein and Dr. Perwitz from the Medical University of Lübeck. We further acknowledge the excellent technical support with the Clariom S array of the Core Facility for DNA technologies of Knut Krohn from the University of Leipzig. For the great assistance in flow-cytometric cell sorting we want to thank the Core Unit Fluorescence Technologies of Kathrin Jäger from the University of Leipzig. We want to thank Claudia Ruffert and Ines Müller for the technical support in the lab and Wenfei Sun and Hua Dong for the support in human AT RNA-sequencing.

## Data availability

Microarray data have been deposited in the ArrayExpress database at EMBL-EBI and are accessible upon publication. The human RNA-seq data from the LOBB have not been deposited in a public repository due to restrictions imposed by patient consent but can be obtained from Matthias Blüher upon request.

