## Supplementary material for "Sex-specific role of epigenetic modification of a leptin upstream enhancer in the adipose tissue": Suppl_Figures

Supplemental Figure 1

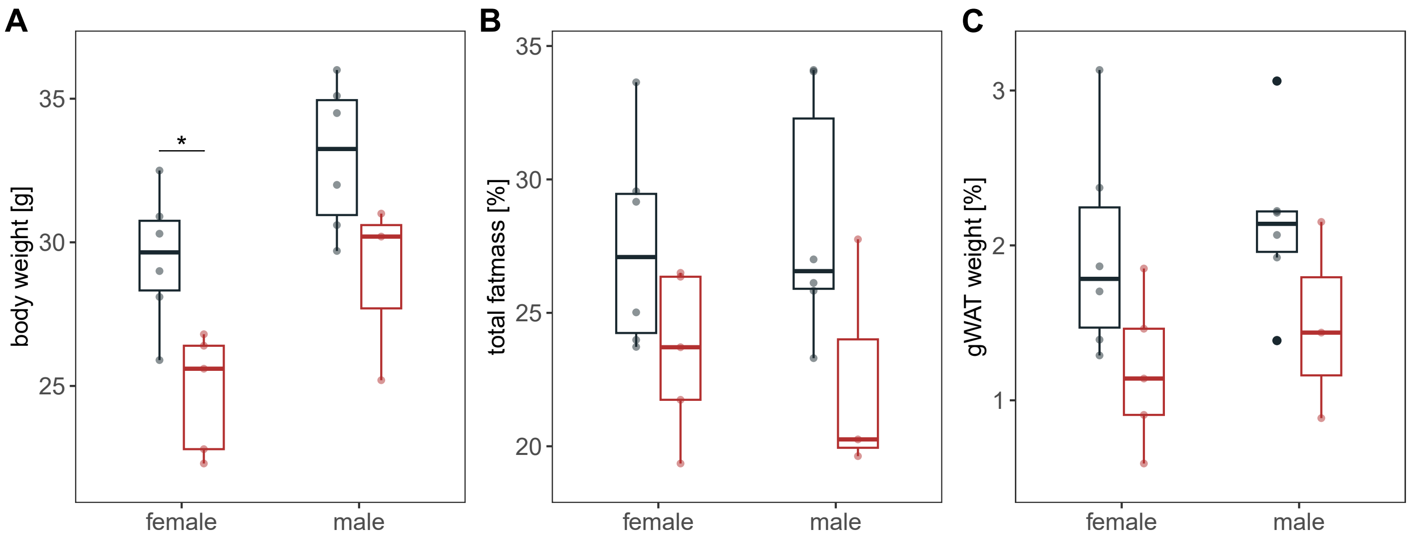


**Supplemental Figure 1 | Differences in offspring's body composition after maternal T3 treatment grouped by sex.** Boxplots show sex specific differences of A) body weight, B) total fatmass (% of body weight), and C) gonadal white adipose tissue mass (gWAT, in % of body weight). Black : offspring of control dams, Red: offspring of T3 treated dams. Significance of differences were calculated by unpaired Wilcoxon rank test. * $P < 0.05$

Supplemental Figure 2

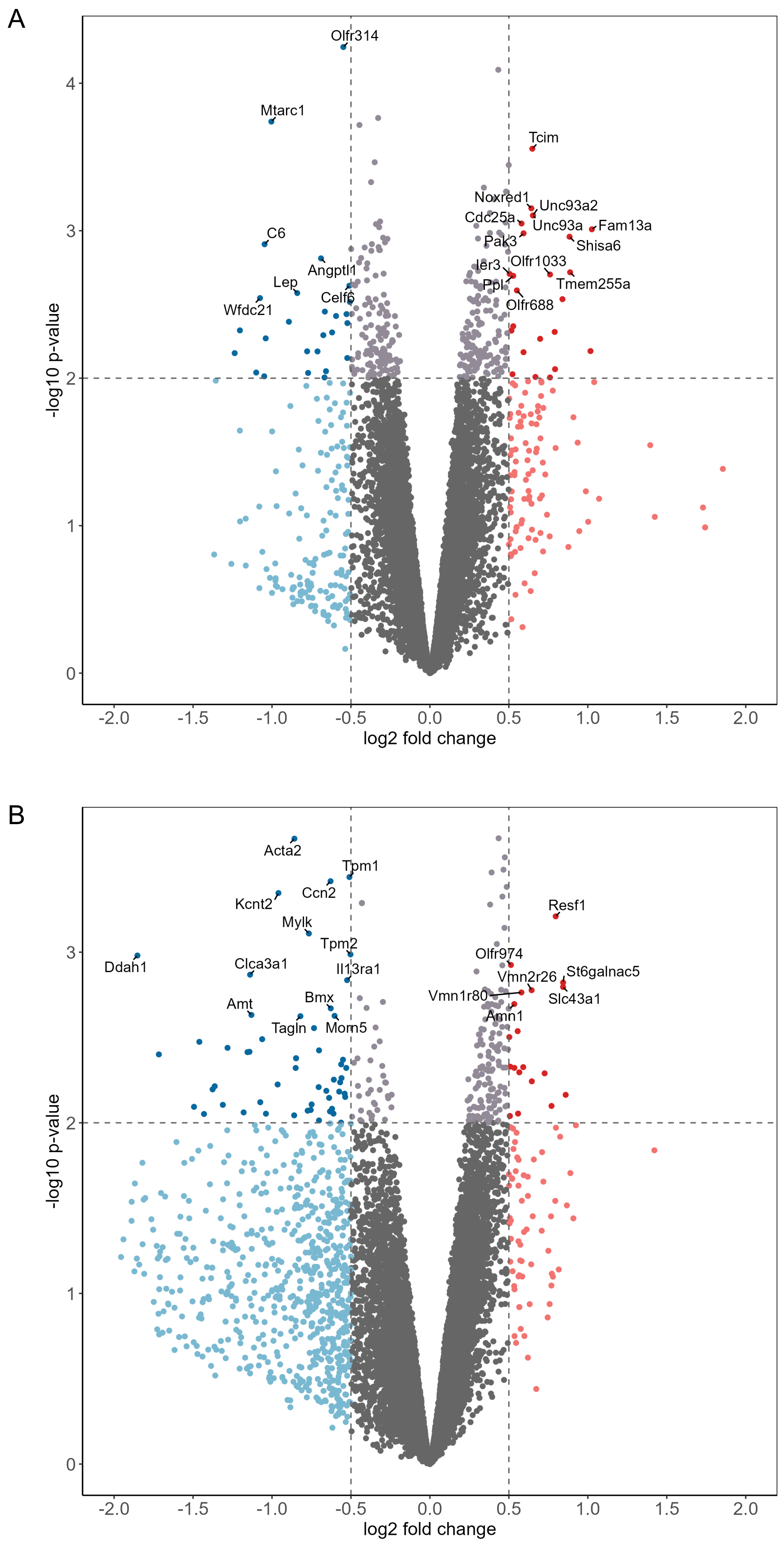


**Suppl Figure 2 | Differentially expressed genes after maternal T3 treatment.**
Volcano Plots showing log 2 fold change of gene expression in gWAT of female offspring (A) and male offspring (B) from T3-treated vs control dams.
Label: top 20 differentially expressed genes with log2 fold change > 0.5 sorted by significance level.
Blue: downregulated genes after maternal T3 treatment, Red: upregulated genes after maternal T3 treatment

Supplemental Figure 3

A

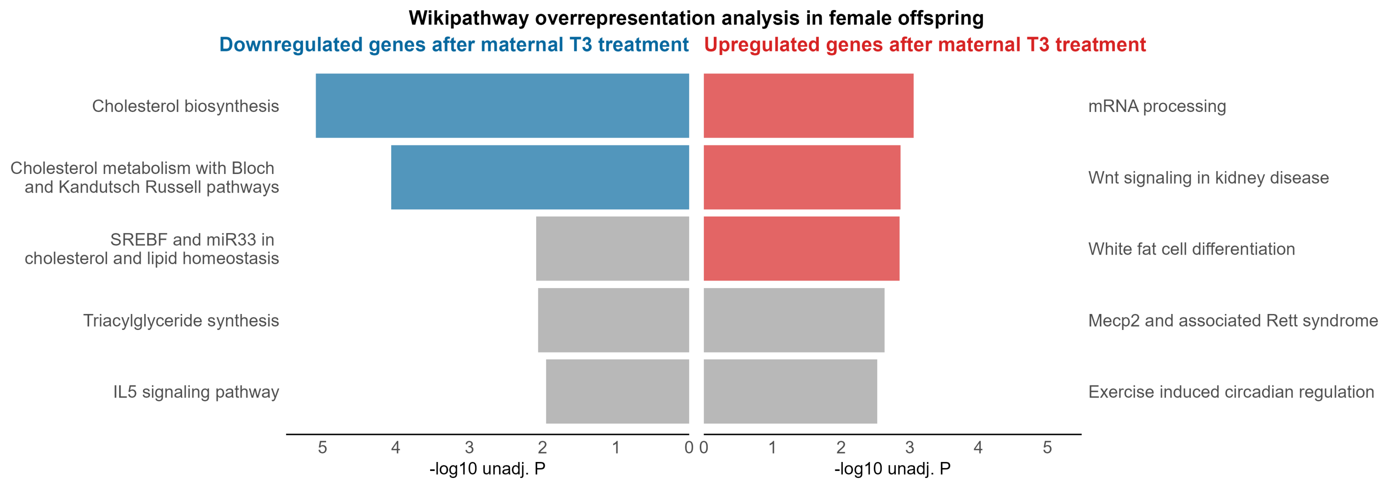


B

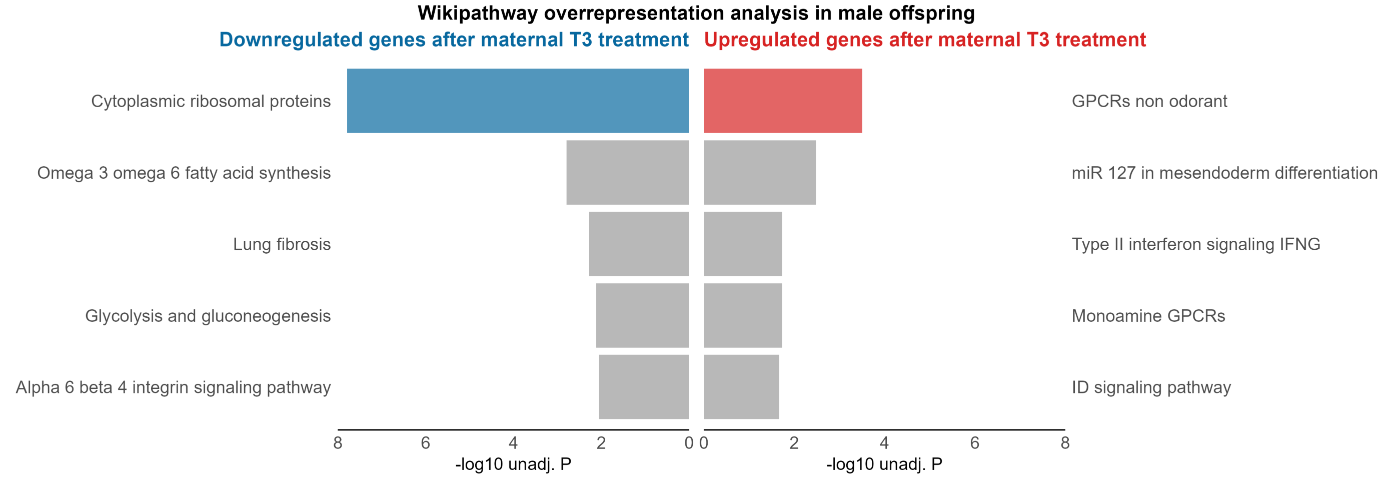


**Suppl Figure 3 | Wikipathway overrepresentation analysis of differentially expressed genes after maternal T3 treatment.** Butterfly plot showing Top 5 overrepresented Wikipathways for differentially expressed genes in gWAT of female offspring (A) and male offspring (B). Blue: Overrepresented pathways of downregulated genes (log2 fold change < 0, unadj. P < 0.05) with an overrepresentation FDR < 0.05;  Red: Overrepresented pathways of upregulated genes (log2 fold change > 0, unadj. P < 0.05) with an overrepresentation FDR < 0.05; Grey:  Overrepresented pathways with an overrepresentation FDR > 0.05, but unadj. overrepresentation P < 0.05

Supplemental Figure 4

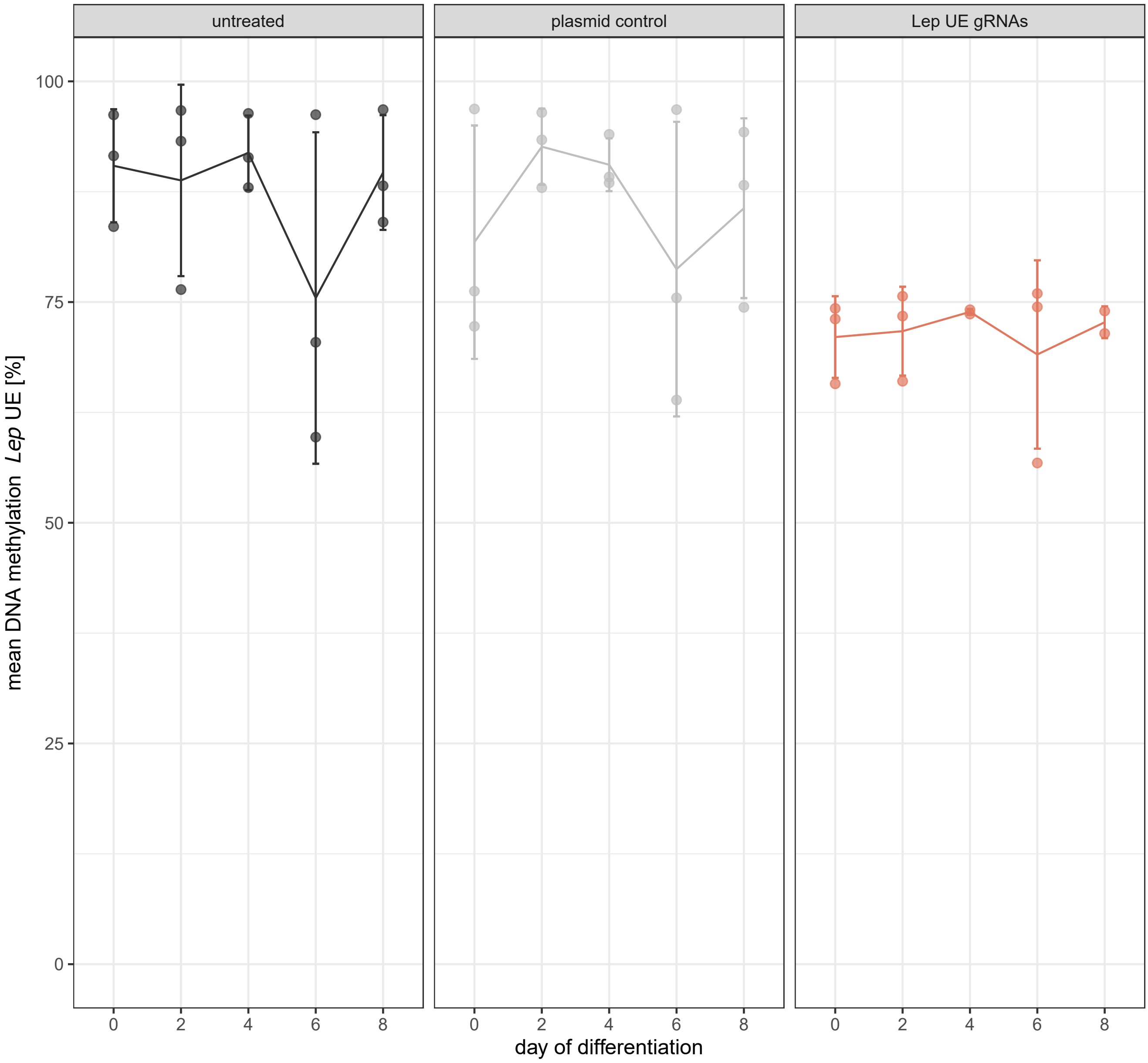


**Supplemental Figure 4 | Course of *Lep* UE DNA methylation during differentiation after demethylation.**
Line plot showing the course of average DNA methylation at the *Lep* upstream enhancer (mean of three CpG sites) in untreated epididymal adipocytes (black), cells transfected with pPlatTET gRNA2 vector without gRNA (plasmid control, gray) and in cells transfected with pPlatTET gRNA2 incl. gRNAs targeting the *Lep* upstream enhancer (Lep UE gRNAs, red). Each dot represents the mean DNA methylation of one out of N=3 experiments. Line connects the mean of all experiments.

### Supplemental Figure 5

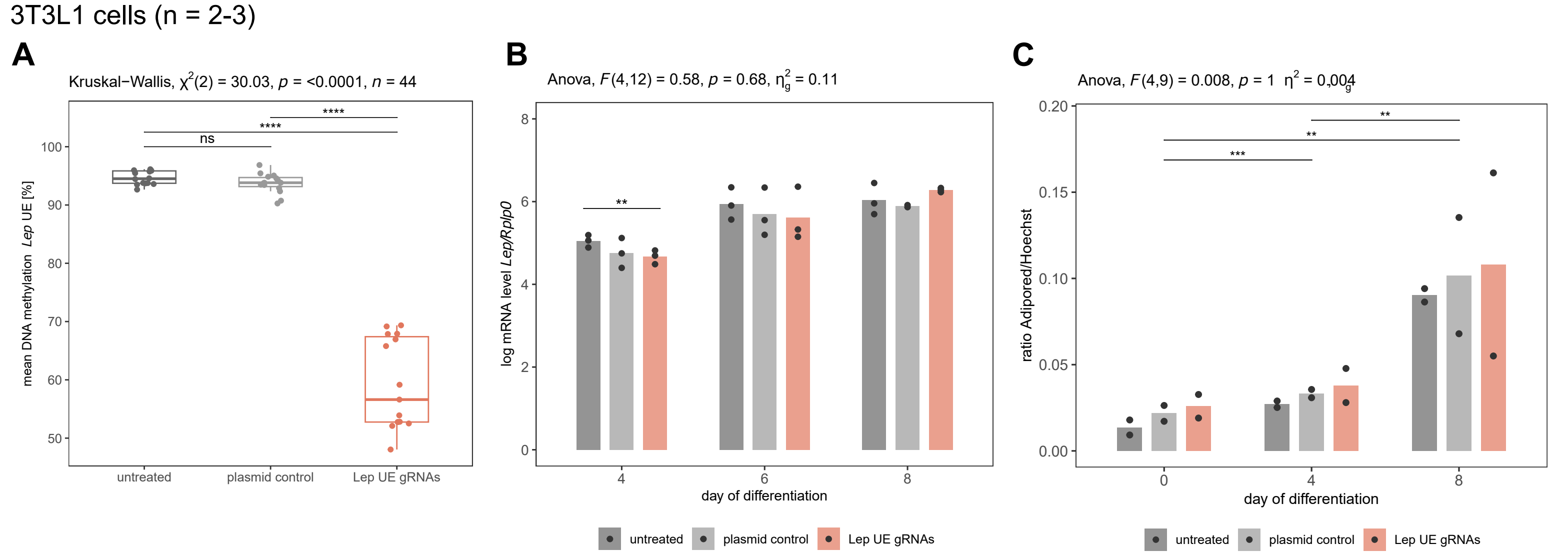


**Supplemental Figure 5 | In vitro Lep UE hypomethylation and effect on adipocyte differentiation in 3T3-L1 cells.**
A) Boxplot shows DNA methylation level [%] of *Lep* UE across all three analysed CpG sites by bisulfite pyrosequencing. Significance of differences between treatments were calculated using Kruskal Wallis test, followed by pairwise comparisons by Wilcoxon rank sum test corrected for multiple testing by FDR (*** FDR > 0.001, ns FDR > 0.05). Dots represent mean methylation levels of day 0, 2, 4, 6 and 8 of differentiation for each experiment (n = 3 experiments x 5 time points).
B) Barplot shows the mean of log transformed *Lep* mRNA levels normalised to *Rplp0* mRNA levels for days 4, 6 and 8 of adipocyte differentiation from n = 3 experiments. Results of mixed two-way ANOVA used to assess the effect of treatment and time on Lep expression are shown on top of the graph. Pairwise comparisons of treatment effects using paired Student's t-test (paired by experiment number) are depicted in the graph. No significant differences were found after correction for multiple testing by FDR. Shown significance level are uncorrected for multiple testing: # P < 0.05.  C) Barplot shows the lipid amounts in cells during differentiation on day 0, 4 and 8 between treatments. Shown is the average ratio of Adipored and Hoechst fluorescence intensity from n = 12 wells in n = 2 experiments. Results of mixed two-way ANOVA used to assess the effect of treatment and time on lipid accumulation are shown on top of the graph. Significance levels from pairwise comparison of time points using paired Student's t-test (paired by experiment number) and corrected for multiple testing by FDR are depicted in the graph (*** FDR < 0.001, ** FDR < 0.01, * FDR < 0.05). No significant differences are found when comparing treatments.

### Supplemental Figure 6

inguinal adipocytes (n =1)

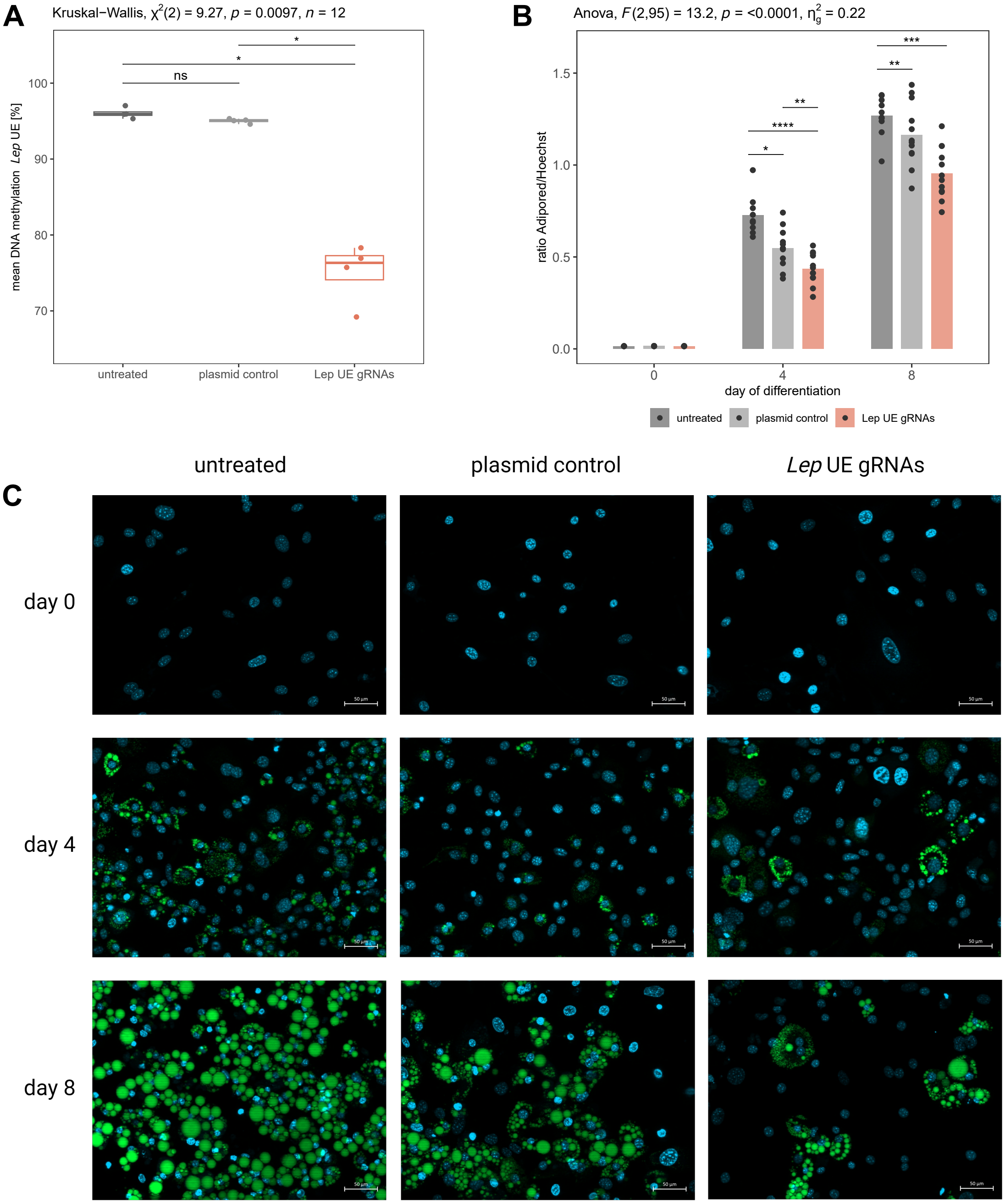


**Supplemental Figure 6 | *In vitro Lep* upstream enhancer (UE) hypomethylation and effect on adipocyte differentiation in female inguinal preadipocytes.**

A) Boxplot shows DNA methylation level [%] of Lep UE across all three analysed CpG sites by bisulfite pyrosequencing. Significance of differences between treatments were calculated using Kruskal Wallis test, followed by pairwise comparisons by Wilcoxon rank sum test corrected for multiple testing by FDR (*** FDR > 0.001, ns FDR > 0.05). Dots represent mean methylation levels of day 0, 4, 6 and 8 of differentiation for each experiment (n = 1 experiment x 4 time points). B) Barplot shows the lipid amounts in cells during differentiation on day 0, 4 and 8 between treatments. Shown is the ratio of Adipored and Hoechst fluorescence intensity from n = 12 wells in n = 1 experiment. Results of mixed two-way ANOVA used to assess the effect of treatment and time on lipid accumulation are shown on top of the graph. Significance levels from pairwise comparison of time points using unpaired Student's t-test and corrected for multiple testing by FDR are depicted in the graph (*** FDR < 0.001, ** FDR < 0.01, * FDR < 0.05). C) Fluorescence microscopy pictures from day 0, 4 and 8 of differentiation in untreated cells, cells transfected with plasmid control and cells transfected with plasmid targeting the Lep UE. Green: lipids stained with Adipored and Blue: DNA stained with Hoechst. Scale bar = 50 µm.
