## Supplementary material for "Sex-specific role of epigenetic modification of a leptin upstream enhancer in the adipose tissue": Suppl_Information

### Supplementary Information

**Adipocyte differentiation**

*Epididymal preadipocytes*

For differentiation FACS sorted cells were seeded into 12 well plates and cultured in growth medium until reaching 100% confluence (usually eight to ten days after electroporation). Adipocyte differentiation was induced 24 h post-confluence (= day 0) by changing to an induction medium consisting of growth medium supplemented with 500 µM isobutylmethylxanthine (IBMX; Sigma Aldrich by Merck, Germany), 2 µg/ml dexamethasone (dexa; ), 0.125 mM indomethacin (indo), 20 nM insulin, and 1 nM triiodothyronine (T3; Sigma Aldrich by Merkc, Germany). From day 2 differentiation cells were grown in differentiation medium (growth medium supplemented with insulin and T3) until day 8 post induction. On day 0, 2, 4, 6, and 8 of differentiation cells were washed with PBS (Gibco, USA), snap frozen in liquid nitrogen and stored at -80 °C until nucleic acid extraction for bisulfite sequencing and real time PCR.

*3T3-L1*

Culturing and transfection of 3T3-L1 cells was done as described for the epididymal preadipocytes. Differentiation was induced 24 h post-confluence by changing to an induction medium consisting of the growth medium supplemented with 500 µM IBMX, 0.2 µg/ml dexa, 1µM insulin. On day 2 the medium is changed to differentiation medium (growth medium supplemented with 1µM insulin) and then changed back to growth medium on day 4 until the end of differentiation.

*Inguinal preadipocytes*

Inguinal preadipocytes were isolated from the inguinal fat pad of female wildtype mice. Immortalisation of the cells was performed by Applied Biological Materials (abm, Canada) by transduction with Lenti-hTERT. Immortalised preadipocytes were grown in DMEM supplemented with 20% FBS and 25 µg/ml ascorbic acid (Sigma Aldrich by Merck, Germany)(=growth medium). Transfection and FACS sorting was performed as described for epididymal preadipocytes. Differentiation was induced 24 h post-confluence by changing to an induction medium consisting of growth medium supplemented with 250 µM IBMX, 0.2 µg/ml dexa, 1µM rosiglitazone, 30 nM insulin, and 1 nM T3. On day 4 the medium is changed back to the growth medium until the end of differentiation.

**Nucleic acid isolation**

Genomic DNA from frozen murine gWAT samples was isolated using the QIAmp DNA Mini Kit (Qiagen, Germany). Tissue samples were cut into small pieces, incubated with Buffer ATL and Proteinase K at 56°C and 650 rpm for 3h, followed by RNA removal with RNase A treatment, and DNA extraction according to manufacturer's protocol. DNA from human AT as well as frozen cell pellets was isolated using the DNeasy blood and tissue kit (Qiagen, Germany) according to the respective manufacturer's instructions for each sample type. DNA sample yield, integrity, and quality were assessed using Quantus™ QuantiFluor dsDNA (Promega, USA) and Nanodrop 2000 spectrometer (Thermo Fisher Scientific, USA).

Total RNA from murine gWAT, as well as human OVAT samples was extracted after homogenization with the Precellys® homogenizer (Bertin Technologies, France) using QIAzol lysis reagent and the RNeasy Lipid Tissue Kit (Qiagen, Germany). RNA from cells of the *in vitro* hypomethylation was extracted after homogenisation with QiaShredder tubes (Qiagen, Germany) and addition of chloroform using the RNeasy Mini Kit (Qiagen, Germany) with on-column DNAse treatment. RNA concentrations and qualities were determined with the Nanodrop 2000 spectrometer.

**RNA sequencing analyses**

To generate ribosomal RNA-depleted RNA sequencing of the human OVAT samples, we followed the SMARTseq protocol (ref). All libraries were sequenced as single-end reads on a Novaseq 6000 instrument (Illumina, USA) at the Functional Genomics Center Zurich, Switzerland. Raw sequencing reads were pre-processed using f*astp* (v0.20.0)^2^ with a minimum read length of 18 nts and a quality cut-off of 20. To align the reads to the reference genome (assembly GRCh38.p13, GENCODE release 32^3^) and quantify gene-level expression, we utilised the pseudoaligner *kallisto^4^*. In cases, where samples had more than 20 million read counts, downsampling was performed to achieve a consistent count of 20 million reads using the R package *ezRun* (v3.14.1; <https://github.com/uzh/ezRun>, accessed on 23 March 2022). Homoscedastic normalisation with respect to library size was performed using the variance-stabilising transformation from DESeq2 (v1.32.0). To account for the impact of in vitro RNA degradation, we adjusted the normalised counts by utilising transcript integrity numbers (TINs) estimated with RSeQC v4.0.0. Analyses were conducted in R v4.3.1 ([www.R-project.org](http://www.r-project.org)).
